# PET microplastics increase the risk of insulin resistance and pancreatitis

**DOI:** 10.1101/2024.10.25.620380

**Authors:** Karol Mierzejewski, Aleksandra Kurzyńska, Monika Golubska, Ismena Gałęcka, Jarosław Całka, Iwona Bogacka

**Author notes:** Aleksandra Kurzyńska –, Monika Golubska –, Jarosław Całka –, Ismena Gałęcka –, Iwona Bogacka –. Corresponding author: Karol Mierzejewski.

## Abstract

Microplastics and their effects on the body have recently been of great concern. Today it is clear that they are not indifferent to human health, but the full spectrum of their impact has not yet been fully described. Pancreatic diseases are becoming increasingly common worldwide, and their etiology is not well understood. Worryingly, these diseases have been increasingly diagnosed in children over the last 20 years, which was previously considered unusual. The aim of the study was therefore to determine the changes in the pancreas caused by PET microplastics in young organisms. For this purpose, the global metabolomic profile of the pancreas of piglets treated with a low (0.1 g/day) or high dose (1 g/day) of PET microplastics for 4 weeks was determined by UPLC-MS analysis. In addition, insulin levels and various biochemical parameters in the blood were analyzed. The study showed that PET microplastics affected the physiological processes in the pancreas at both low and high doses. We found that PET microplastics increased the tissue levels of important metabolites such as glucose, γ-aminobutyric acid, lysophosphatidylcholine or lysophosphatidylethanolamine. In addition, PET microplastics increased blood insulin concentrations and dose-dependently regulated lipase, cholesterol and calcium levels. These results suggest that PET microplastics increase the risk of insulin resistance and pancreatitis.

## 1. Introduction

Microplastics (MPs), defined as plastic particles smaller than 5 mm, have become ubiquitous in the environment due to widespread plastics production and inadequate waste management. One of the most widely used plastics is polyethylene terephthalate (PET) and is found in a variety of consumer products such as drinks bottles, food packaging and textiles. The pervasive presence of microplastics in ecosystems, food and beverages raises major concerns about the potential impact on human health, particularly through ingestion and subsequent accumulation in various organs. The accumulation of MPs has already been demonstrated in many different tissues including brain, liver, placenta, testis or blood (Horvatits et al., 2022; Hu et al., 2024; Ragusa et al., 2021). Plastic particles have been shown to induce neurotoxicity, impair the synthesis of steroid hormones, impact the quality of oocytes and sperm, promote cardiovascular disease, develop microbial dysbiosis as well as hepatic inflammation(Ullah et al., 2023). There is also growing body of evidence that microplastics impair the pancreatic functions.

The pancreas is a vital organ that fulfils both endocrine and exocrine functions that are essential for the proper functioning of the organism. The central role of the pancreas is to regulate macronutrient digestion and thus metabolism/energy homeostasis through the release of various digestive enzymes and pancreatic hormones (Leung, 2010). It consists of two structurally and functionally integrated glandular systems, namely the exocrine and endocrine pancreas. The largest part of this organ consists of acinar or exocrine cells that secrete pancreatic juice, which contains digestive enzymes such as amylase, pancreatic lipase and trypsinogen, into the pancreatic and the accessory pancreatic duct (Röder et al., 2016). The endocrine pancreas, on the other hand, consists of the islets of Langerhans, which secrete hormones such as insulin directly into the bloodstream to regulate blood glucose levels(Al-Suhaimi et al., 2022). Dysfunction of the pancreas can lead to serious diseases such as insulin resistance, diabetes, chronic pancreatitis and pancreatic cancer.

There is evidence that polystyrene MPs exacerbate pancreatic injury and inflammation in acute pancreatitis in mice (Zheng et al., 2023). In addition, exposure to polystyrene MPs induces insulin resistance in mice, which is mediated by regulating the gut microbiota, stimulating inflammation and inhibiting the insulin signaling pathway (Huang et al., 2022). Another study showed that MPs induce oxidative stress and activate the GRP78/CHOP/Bcl-2 signaling pathway to increase pancreatic apoptosis in mice (Wang et al., 2022). In our previous study, we found that PET MPs in piglets alter the expression of miRNA in serum-derived extracellular vesicles (EVs), which are associated with insulin resistance, type II diabetes and the development of pancreatic cancer (Mierzejewski et al., 2023). The available literature on the effects of microplastics on the pancreas is patchy and a comprehensive understanding of the mechanisms is still lacking. Therefore, the aim of this study was to investigate the effects of oral treatment of piglets with PET MPs on the global metabolomic profile of the pancreas using UPLC-MS/MS analysis. In addition, insulin levels and various biochemical parameters in the blood were determined.

## 2. Materials and methods

### 2.1. Animals

All experimental protocols were approved by the Local Ethics Committee of the University of Warmia and Mazury in Olsztyn (Decision No. 10/2020 of 26 February 2020) and the study was conducted in accordance with the provisions of the European Union Directive on the ethical use of experimental animals (EU Directive 2010/63/EU for animal experiments). All animals were housed in breeding pens under standard laboratory conditions and were provided with free access to fresh water (ad libitum) and an age-appropriate feed mixture. The temperature in the animal pens was maintained at 20-22°C and the humidity was between 55% and 60%. All plastic objects were removed from the animals’ environment. The watering through and feeders in the experimental pens were made of stainless steel. The equipment on the farm where the gilts were reared before the experiment was also made of stainless steel. The elements of the environment that guaranteed a high level of welfare were made of wood or bedding. The experiment lasted 4 weeks and was carried out on 8-week-old (Pietrain x Duroc) immature gilts (n = 15) with an estimated body weight of 20 kg. The animals were divided into three groups: 1) control group (CTR; n = 5), which received empty gelatine capsules per os; 2) experimental group (LD; n = 5), which received a low dose of PET MPs (0.1 g/pig/day in gelatine capsules) per os; 3) experimental group (HD; n = 5), which received a high dose of PET MPs (1 g/pig/day in gelatine capsules) per os. The gelatine capsules were administered to the gilts 1 hour before morning feeding. The plastic used for the experiment is a semi-crystalline polyethylene terephthalate powder (Goodfellow Cambridge Ltd., England). For particle analysis, 500 randomly selected particles were analyzed microscopically using the Zeiss Axio Imager.M2 fluorescence microscope (Zeiss, Germany) and ZEISS ZEN Microscopy Software (Zeiss, Germany). Randomization was performed by taking a representative sample of the whole material from different depth levels and then dividing it into 25 slides. Then 20 randomly selected particles were measured on each slide. The average length of the larger side of the molecule was 153.09 μm, MIN – 1.25 μm, MAX – 299.75 μm, SD – 85.13, SEM – 3.81. Different shapes of the particles (spherical, fibrous, irregular) were observed. The PET particles had both sharp and rounded edges (Ismena Gałęcka and Jarsoław Całka, 2024). The doses administered were selected based on the weekly human consumption of MPs reported in the literature (Deng et al., 2017; Senathirajah et al., 2021) and adjusted to the weight of the gilts. After 4 weeks, the piglets were euthanized. The euthanasia protocol was based on the use of atropine (0.05 mg/kg i.m., Polfa, Poland), followed by xylazine (3 mg/kg i.m., Vet-Agro, Poland) and ketamine (6 mg/kg i.m., Vetoquinol Biowet, Poland). After approximately 20 min, when the gilts were unconscious, an overdose of sodium pentobarbital (0.6 mL/kg i.v., Biowet, Poland) was applied (Gałęcka et al., 2024). Once it was confirmed that vital functions had ceased (absence of pupillary reflex, pulse and respiration), the material was immediately collected for further analyses.

### 2.2. Sample preparation

The sample was thawed on ice, approximately 100 mg of sample was weighed into a tube, 80% methanol containing 8 μL/mg of the raw material and two 5 mm metal spheres were added to the tube. All samples were ground twice for 180 s at 65 Hz, followed by sonication for 30 min at 4°C. Then each sample was kept at -20°C for 1 hour. Afterwards, the samples were centrifuged for 15 min at 12,000 rpm at 4°C. Finally, 200 μL of the supernatant and 5 μL of DL-o-chlorophenylalanine (0.2 mg/mL) were transferred to a vial for LC-MS analysis. Quality control (QC) samples were used to evaluate the methodology. The same amount of extract was obtained from each sample and mixed as QC samples. The QC sample was prepared using the same sample preparation procedure (Polak et al., 2023).

### 2.3. UPLC-MS

Separation of compounds was performed by ultra-performance liquid chromatography (UPLC) coupled with tandem mass spectrometry using a Q Exactive Plus MS (Thermo Fisher Scientific, Waltham, MA, United States) and screened with electrospray ionization mass spectrometry (ESI-MS). The LC system comprised an ACQUITY UPLC HSS T3 column (100 × 2.1 mm, 1.8 μm) (Waters Corporation, Milford, MA, United States). The mobile phase was composed of solvent A (0.05% formic acid in water) and solvent B (acetonitrile) with a gradient elution (0–1 min, 5% B; 1–12 min, 5%–95% B; 12–13.5 min, 95% B; 13.5–13.6 min, 95%–5% B; 13.6–16 min, 5% B). The flow rate of the mobile phase was 0.3 mL/min. The column temperature was maintained at 40°C, and the sample manager temperature was set at 4°C. Mass spectrometry parameters in ESI+ and ESI-modes are listed as follows: for ESI+ mode, the heater temperature was 300°C, the sheath gas flow rate was 45 arbitrary units (arb), the auxiliary gas flow rate was 15 arb, the sweep gas flow rate was 1 arb, the spray voltage was 3.0 kV, the capillary temperature was 350°C, and the S-Lens RF level was 30%. For ESI-mode, the heater temperature was 300°C, the sheath gas flow rate was 45 arb, the auxiliary gas flow rate was 15 arb, the sweep gas flow rate was 1 arb, the spray voltage was 3.2 kV, the capillary temperature was 350°C, and the S-Lens RF level was 60%. The mass spectrometry scan modes included Full Scan (m/z 70–1050, resolution: 70,000) and data-dependent MS2 (dd-MS2, TopN = 10, resolution: 17,500) with higher-energy collisional dissociation (HCD) as the collision mode.

### 2.4. Data processing and statistical analysis

Raw LC-MS/MS data were acquired and aligned using the Compound Discoverer (v. 3.0, Thermo Fisher Scientific, Waltham, MA, United States) based on the m/z value and retention time (RT) of the ion signals. The ions from the ESI and ESI+ modes were merged and imported into SIMCA-P software (version 14.1, Sartorius, Göttingen, Germany) for multivariate analysis. The spectral data were normalized and automatically scaled prior to statistical analysis. Principal component analysis (PCA) was initially used as an unsupervised method to visualise the data and identify outliers. Supervised regression modelling was performed on the dataset using Partial Least Squares Discriminant Analysis (PLS-DA) or Orthogonal Partial Least Squares Discriminant Analysis (OPLS-DA) to identify potential biomarkers. The quality of the models was assessed using the relevant R2 and Q2 values, where R2 indicates the variance explained in the model and Q2 reflects the predictability of the model (Fig. 1A-B, 2A-B). The importance of each ion in the PLS-DA or OPLS-DA models was assessed using the VIP (Variable Importance in the Projection) score, where VIP values greater than 1.5 were considered significant (Fig. 1C, Fig. 2C, Fig. 3A, Fig 4A). Biomarkers were filtered and confirmed by combining the results of the VIP scores and the t-test (p < 0.05). The chemical structures and IDs of the metabolites were identified using the Human Metabolome Database (Wishart et al., 2012), the KEGG Database (Kanehisa, 2000; Kanehisa et al., 2016), the PubChem Compound ID Database (Kim et al., 2022) and the ChemSpider Database (https://www.chemspider.com/Default.aspx). In the multivariate analyses, hierarchical cluster analysis (HCA) with the measured Euclidean distance and the average cluster algorithm was used to visualize the differences in the concentration of each statistically significant metabolite between the groups in two different ESI modes. Metabolites were then assigned to substance groups and pathway enrichment analysis was performed using MetaboAnalyst 5.0 (Pang et al., 2021; Xia et al., 2009).

**Figure 1.**
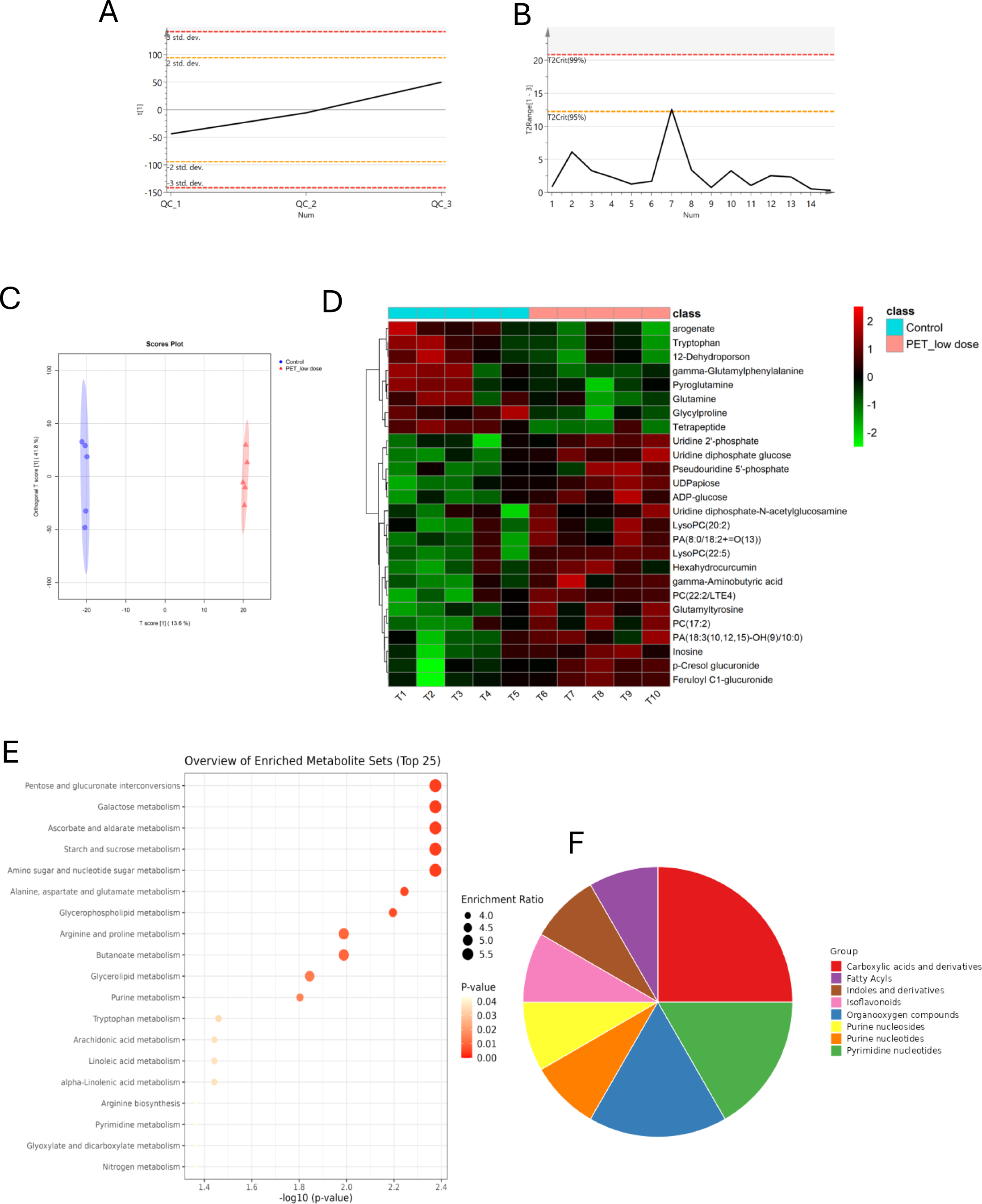
The analysis of untargeted metabolome profiles of the pancreas isolated from the piglets treated with a low dose of PET microplastics (0.1 g/day) compared to the control group in negative ionization mode (ESI-) **(A)** The PCA score plot of the QC samples in ESI-; the X axis indicates the number of QC samples, and the Y axis indicates the range of RSD. **(B)** The line plot of the normalized QC samples in ESI-mode; the X axis indicates the number of QC samples, and the Y axis indicates the 95% confidence interval. **(C)** The scatter plots of the OPLS-DA model. **(D)** Hierarchical cluster analysis of biomarkers. Color intensity correlates with degree of increase (red) and decrease (green) relative to the mean metabolite ratio. **(E)** Dot plot of network analysis in differentially regulated metabolites according to KEEG and MataboAnalyst. **(F)** Distribution of differentially regulated metabolites by different metabolite classes identified in ESI-mode.

**Figure 2.**
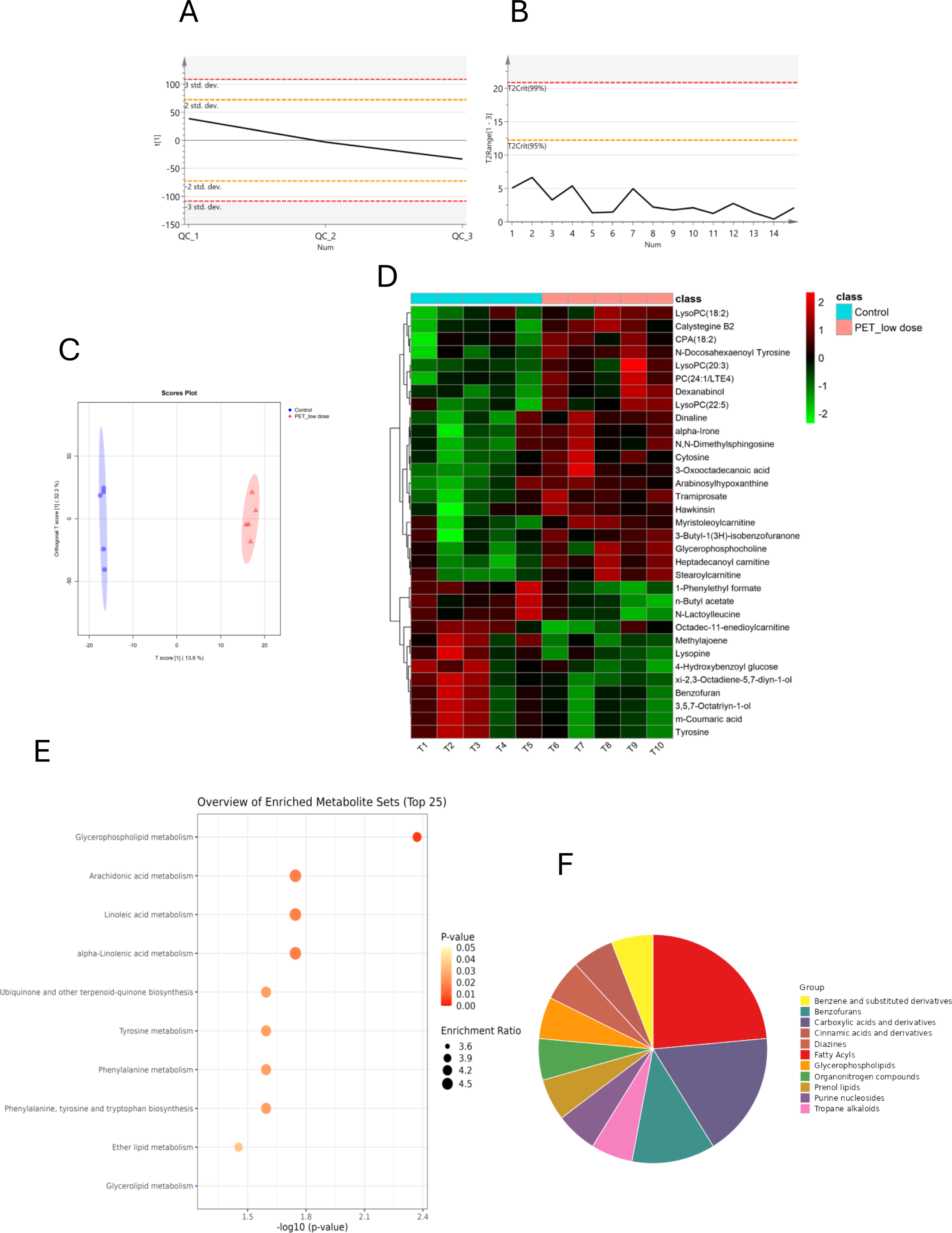
The analysis of untargeted metabolome profiles of the pancreas isolated from the piglets treated with a low dose of PET microplastics (0.1 g//day) compared to the control group in positive ionization mode (ESI+) **(A)** The PCA score plot of the QC samples in ESI+; the X axis indicates the number of QC samples, and the Y axis indicates the range of RSD. **(B)** The line plot of the normalized QC samples in ESI+ mode; the X axis indicates the number of QC samples, and the Y axis indicates the 95% confidence interval. **(C)** The scatter plots of the OPLS-DA model. **(D)** Hierarchical cluster analysis of biomarkers. Color intensity correlates with degree of increase (red) and decrease (green) relative to the mean metabolite ratio. **(E)** Dot plot of network analysis in differentially regulated metabolites according to KEEG and MataboAnalyst. **(F)** Distribution of differentially regulated metabolites by different metabolite classes identified in ESI+ mode.

**Figure 3.**
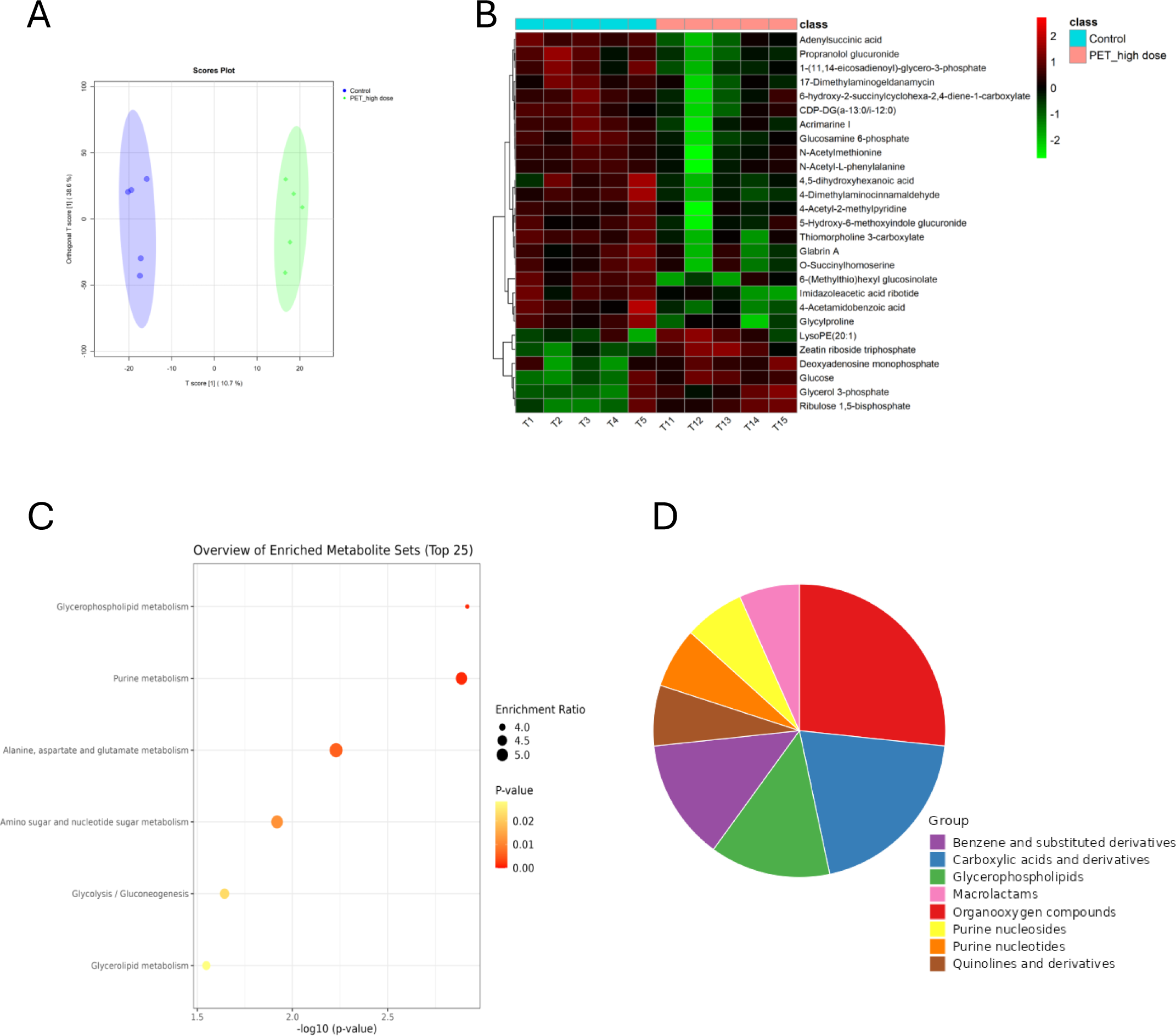
The analysis of untargeted metabolome profiles of the pancreas isolated from the piglets treated with a high dose of PET microplastics (1 g/day) compared to the control group in negative ionization mode (ESI-). **(A)** The scatter plots of the OPLS-DA model. **(B)** Distribution of differentially regulated metabolites by different metabolite classes identified in ESI-mode. **(C)** Hierarchical cluster analysis of biomarkers. Color intensity correlates with degree of increase (red) and decrease (green) relative to the mean metabolite ratio. **(D)** Dot plot of network analysis in differentially regulated metabolites according to KEEG and MataboAnalyst.

**Figure 4.**
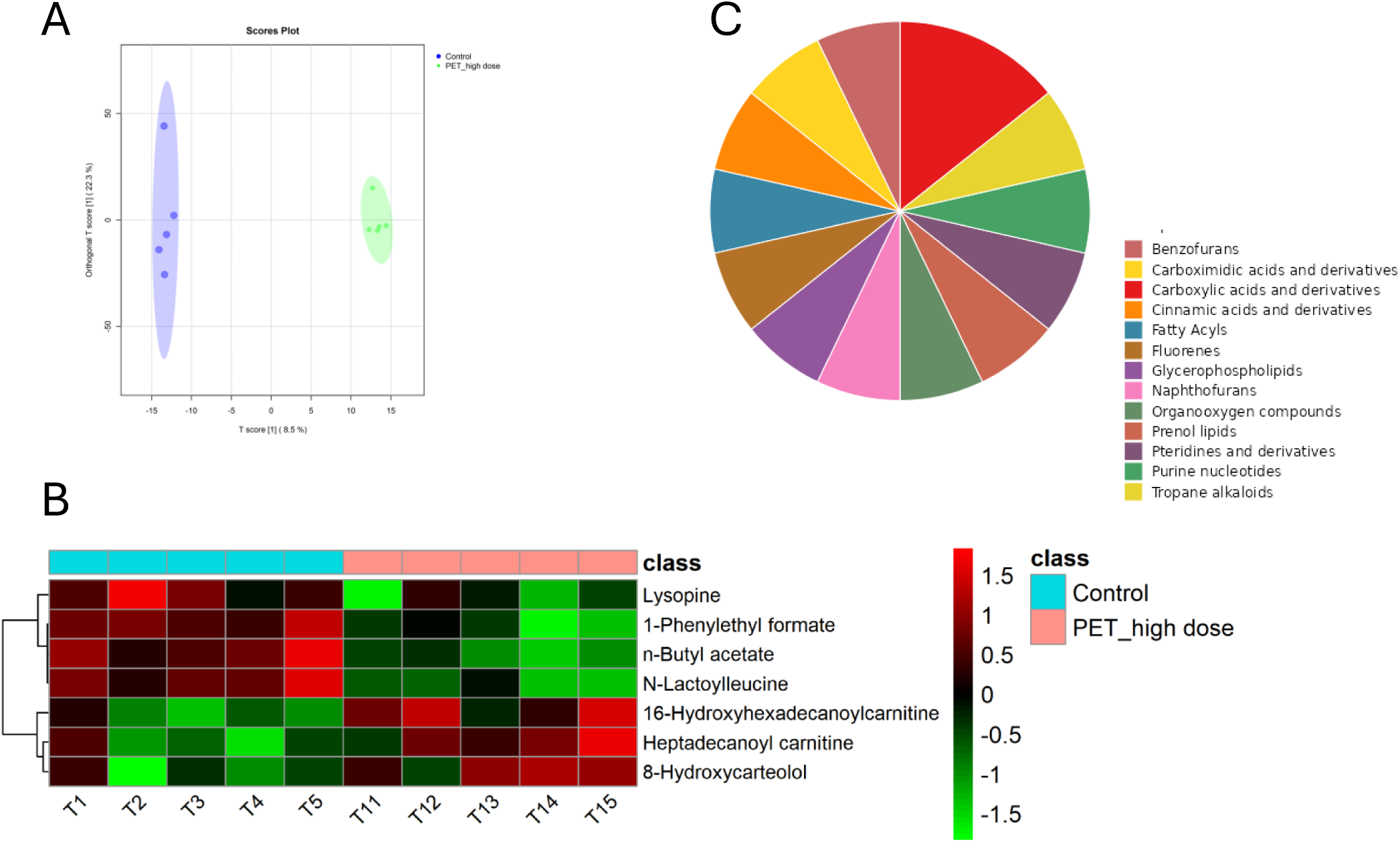
The analysis of untargeted metabolome profiles of the pancreas isolated from the piglets treated with a high dose of PET microplastics (1 g/day) compared to the control group in positive ionization mode (ESI+). **(A)** The scatter plots of the OPLS-DA model. **(B)** Distribution of differentially regulated metabolites by different metabolite classes identified in ESI+ mode. **(C)** Hierarchical cluster analysis of biomarkers. Color intensity correlates with degree of increase (red) and decrease (green) relative to the mean metabolite ratio.

### 2.5. Biochemical analysis

Prior to blood sampling, the pigs were subjected to a 24-hour fasting period and had ad libitum access to water. Blood was collected by puncturing the external jugular vein into a tube containing a coagulation activator. It was then left to stand at room temperature for at least 30 min. To obtain the serum, the collected material was centrifuged at room temperature for 10 min at 3,000 rpm. The collected serum was aliquoted and stored at -20°C for biochemical analyses. The biochemical analysis included the determination of the concentration of albumin (ALB), alkaline phosphatase (ALKP), alanine aminotransferase (ALT), amylase (AMY), blood urea nitrogen (BUN), calcium (Ca), cholesterol (CHOL), creatinine (CREA), fructosamine (FRU), gamma-glutamyl transferase (GGT), globulins (GLOB), glucose (GLU), lactate dehydrogenase (LDH), lipase (LIPA), phosphorus (PHOS), total bilirubin (TBIL) and total protein (TP) using the Idexx Catalyst One Chemistry Analyser (IDEXX Laboratories, Inc., United States) according to the manufacturer’s instructions. Based on the results obtained, the analyser also calculated the ALB/GLB and BUN/CREA ratios. For parameters whose concentration did not exceed the minimum value detectable by the analyser (FRU 100 µmol/l, TBIL 0.1 mg/dl, LIPA 10 U/l), the minimum value of the assay (FRU 100 µmol/l, TBIL 0.1 mg/dl, LIPA 10 U/l) was assumed for further analysis. The concentration of insulin in the serum was determined using a commercially available ELISA kit (ELK Biotechnology, cat: ELK1385) according to the manufacturer’s protocol and as previously described (Mierzejewski et al., 2022). The range of standard curves was 7.82-500 pg/mL. Absorbance values were measured at 450 nm using Infinite M200 Pro reader with the Tecan i-control software (Tecan, Switzerland). The intra- and inter-assay coefficients of variation of the ELISA assay for insulin were < 8% and <10%, respectively. Then, the quantitative insulin sensitivity check index (QUICKI) was calculated, as indicated:

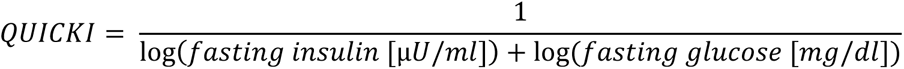

The results obtained were analyzed with Statistica 13.3 software (TIBCO Software Inc., Palo Alto, USA), using one-way analysis of variation (ANOVA) with the Dunnett test. The results obtained were presented as mean ± standard deviation. The results were considered statistically significant at a p-value<0.05 (*p<0.05, **p<0.01). GraphPad Prism 9.0.0 software (Boston, USA) was used for visualization.

## 3. Results

### 3.1. Effect of a low dose of PET microplastics on global metabolomic profile of the pancreas

In the ESI-mode, our analysis revealed 26 biomarkers – 8 of them were downregulated, while 18 were upregulated (Fig. 2D). Most of these differentially regulated metabolites belong to the following classes: carboxylic acids and derivatives (gamma-aminobutyric acid; gamma-glutamylphenylalanine; glutamine; glycylproline; glutamyltyrosine; pyroglutamine), organooxygen compounds (pseudouridine 5’-phosphate; uridine 2’-phosphate; p-cresol glucuronide) or pyrimidine nucleotides (uridine diphosphate glucose; uridine diphosphate-N-acetylglucosamine; UDP-D-apiose) (Fig. 1F). According to the KEGG analysis, the differentially regulated metabolites are involved in processes such as pentose and glucuronate interconversions, galactose metabolism, starch and sucrose metabolism, glycerolipid metabolism or arachidonic acid metabolism (Fig. 1E). Detailed results of the normalized data, HCA and KEGG analysis are presented in the Supplemental Materials (Table S1 a-c).

In ESI+ mode, the analysis revealed 33 biomarkers – 12 of them was downregulated, while 21 were upregulated (Fig. 2D). Most of these differentially regulated metabolites belong to the following classes: fatty acyls (stearoylcarnitine; heptadecanoyl carnitine; CPA(18:2(9Z,12Z)/0:0); 3-oxooctadecanoic acid; 3,5,7-octatriyn-1-ol; xi-2,3-octadiene-5,7-diyn-1-ol; myristoleoylcarnitine); amino acids, peptides, and analogues (N-lactoylleucine, lysopine, hawkinsin), carboxylic acids and derivatives (L-tyrosine; hawkinsin; n-butyl acetate; L-lysopine; N-lactoylleucineor) or benzofurans ((S)-3-butyl-1(3H)-isobenzofuranone; benzofuran) (Fig. 2F). According to the KEGG analysis, the differentially regulated metabolites are involved in processes such as glycerophospholipid metabolism, arachidonic acid metabolism, linoleic acid metabolism or glycerolipid metabolism (Fig. 2E). Detailed results of the normalized data, HCA and KEGG analysis are presented in the Supplemental Materials (Table S2 a-c).

### 3.2. Effect of a high dose of PET microplastics on global metabolomic profile of the pancreas

In the ESI-mode, we identified 27 biomarkers – 21 of them were downregulated, while 6 were upregulated (Fig. 3B). Most of these differentially regulated metabolites belong to the following classes: organooxygen compounds (D-glucose; glucosamine 6-phosphate; 5-hydroxy-6-methoxyindole glucuronide; 6-(methylthio)hexyl glucosinolate; 4-acetyl-2-methylpyridine), carboxylic acids and derivatives (N-acetyl-L-phenylalanine; glycylproline; N-acetyl-L-methionine; thiomorpholine 3-carboxylate) or glycerophospholipids (glycerol 3-phosphate; 1-(11Z,14Z-eicosadienoyl)-glycero-3-phosphate) (Fig. 3D). According to the KEGG analysis, the differentially regulated metabolites are involved in processes such as glycerophospholipid metabolism, purine metabolism, alanine, aspartate and glutamate metabolism, amino sugar and nucleotide sugar metabolism, glycolysis/gluconeogenesis and glycerolipid metabolism (Fig. 3C). Detailed results of the normalized data, HCA and KEGG analysis are presented in the Supplemental Materials (Table S3 a-c).

In ESI+ mode, the analysis revealed 7 biomarkers – 4 of them was downregulated, while 3 were upregulated (Fig. 4C). The differentially regulated metabolites belong to the following classes: carboxylic acids and derivatives (creatinine; L-glutamic gamma-semialdehyde; proline betaine), pteridines and derivatives (riboflavin) or fatty acyls (5-oxo-12-HETE) (Fig. 4B). These differentially regulated metabolites were not assigned to any KEGG pathway. Detailed results of the normalized data, HCA and KEGG analysis are presented in the Supplemental Materials (Table S4 a-c).

### 3.3. Effect of PET microplastics on insulin and blood biochemical parameters

Serum insulin concentrations were higher in piglets treated with either a low or high dose of PET microplastics than in control subjects (Fig. 5). In addition, serum cholesterol levels (CHOL) increased after treatment with a low dose of PET microplastics, while a high dose of PET increased pancreatic lipase (LIPA) levels and decreased calcium (Ca) levels. Other biochemical parameters in serum were not affected by PET microplastics (Fig. 4). The quantitative insulin sensitivity check index (QUICKI) was significantly reduced in the presence of a low dose, while it did not change after a high dose of PET microplastics (Fig. 5).

**Figure 5.**
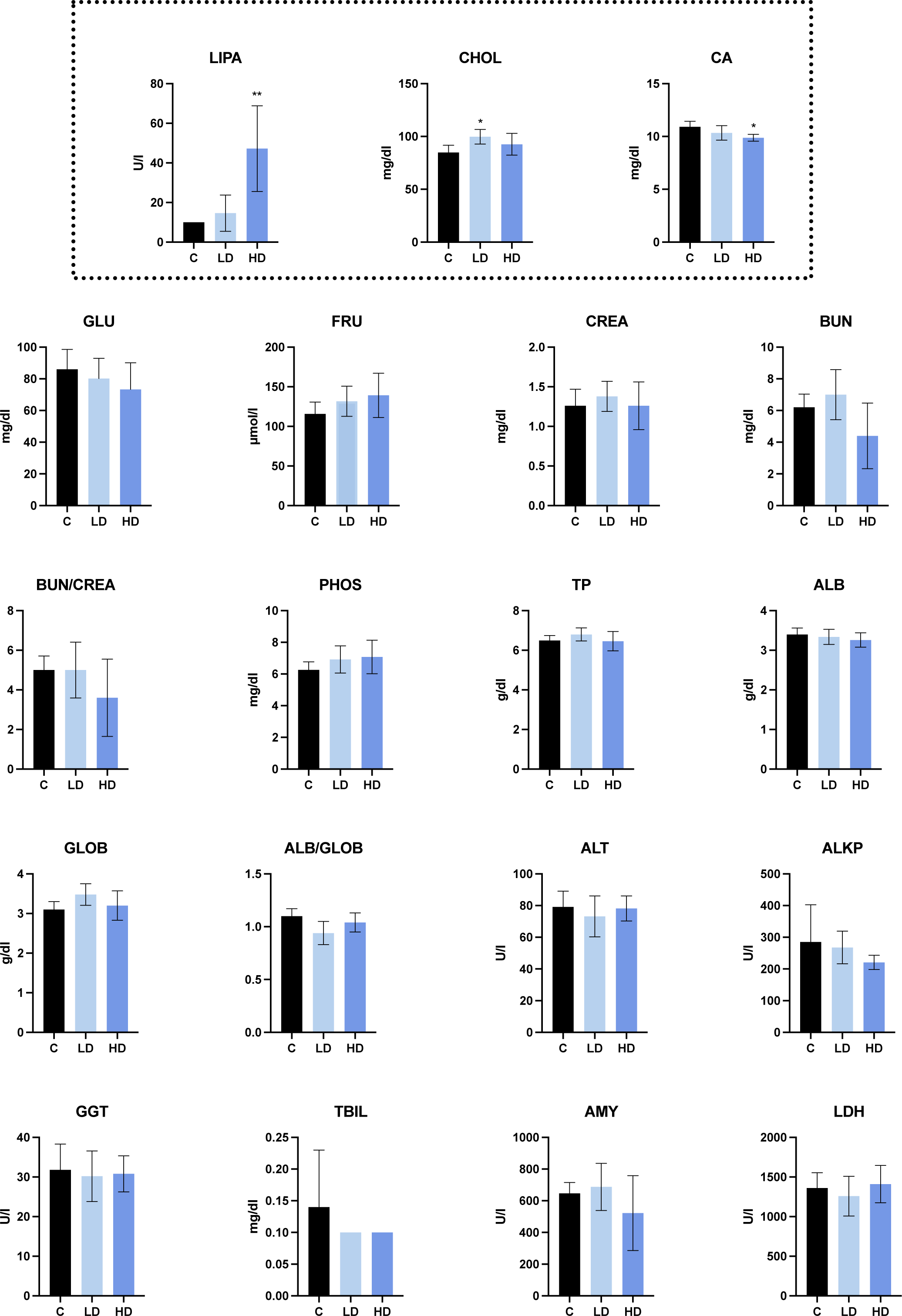
Biochemical parameters measured in the blood of piglets treated with low (LD) and high (HD) doses of PET microplastics, compared to a control group (C).

**Figure 6.**
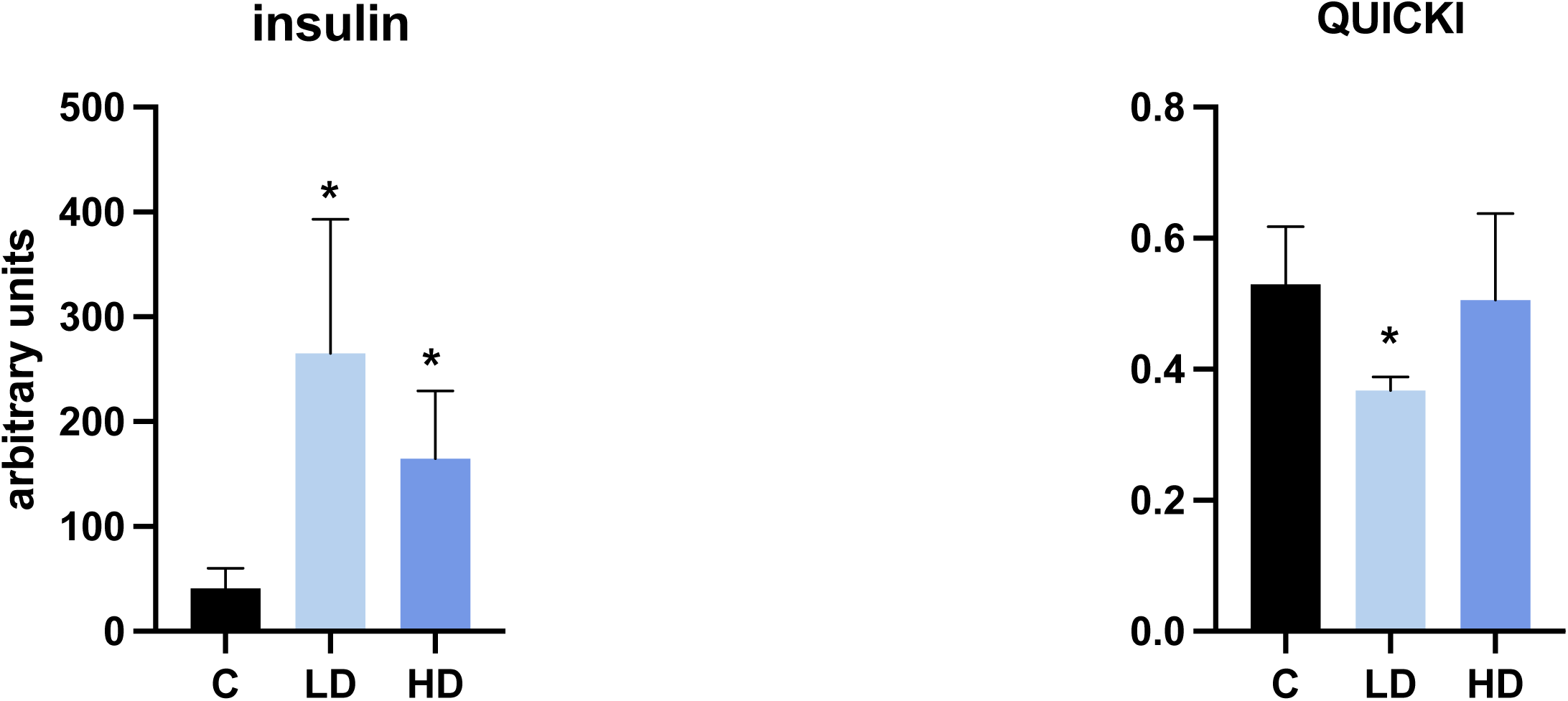
Blood insulin levels and QUICKI parameter in piglets treated with low (LD) and high (HD) doses of PET microplastics, compared to a control group (C).

## 4. Discussion

Pancreatic diseases are an increasing concern in highly developed countries. Similarly, microplastics have become a significant global problem, as there is growing evidence that they contribute to various disorders. Therefore, we investigated the effects of microplastics on the pancreatic metabolome to better understand their potential role in exacerbating pancreatic diseases. In this study, we demonstrated that PET microplastics alter the profile of metabolites in the pancreas and that dysregulation is associated with insulin resistance, glucose and lipid homeostasis and oxidative stress.

One of the most important functions of the pancreas is to regulate glucose homeostasis by releasing glucagon from the α-cells during hypoglycemia and insulin from the β-cells when glucose levels increase (Röder et al., 2016). Abnormal functioning of these cells can lead to impaired glycemic control, which can result in the development of diabetes (Dunning et al., 2005). Type 2 diabetes is characterized by a combination of insulin resistance and changes in β-cell function. Several studies have shown that the β-cell mass in type 2 diabetes is ∼50% of normal (Clark et al., 1988; WOLFFENBUTTEL and HAEFTEN, 1993) and the changes are associated with hyperglycemia (Donath et al., 1999; Eizirik, 1996). High glucose levels in human islet cells alter the balance between pro- and anti-apoptotic BCL proteins towards apoptosis, favoring β-cell death (Federici et al., 2001). Moreover, chronic in vitro exposure to elevated glucose levels renders β-cells hypersensitive to glucose, reducing the threshold for insulin secretion (Jonas et al., 2009; Khaldi et al., 2004; Prentki et al., 2020). There is also evidence that high glucose concentrations stimulate the activation of rat pancreatic stellate cells (PSCs) via the PKC-p38 MAP kinase pathway, suggesting that high glucose may exacerbate pancreatic fibrosis (Nomiyama et al., 2007). The present study shows an increased glucose level in the pancreas of piglets fed a high dose of PET microplastics. Interestingly, blood glucose levels were not significantly altered by PET microplastics, which could indicate an increased uptake of glucose by pancreatic cells under these conditions. Therefore, we hypothesized that elevated glucose levels in the pancreas caused by PET microplastics could affect the proper function of β-cells through various mechanisms such as apoptosis or inflammation.

Our study has shown that both doses of PET microplastics increase serum insulin levels. In healthy individuals, insulin binds to cell surface receptors and triggers responses that enable the uptake of glucose from the blood. In the case of insulin resistance, the cells respond inadequately to insulin, reducing glucose absorption and raising blood glucose levels. To compensate for this, the pancreas increases insulin production, leading to hyperinsulinemia. Impaired insulin secretion in metabolic disorders can precede and worsen insulin resistance (Thomas et al., 2019). Our analysis revealed that the quantitative insulin sensitivity check index (QUICKI) was significantly lower in piglets treated with a low dose of PET microplastics, suggesting an increased risk of insulin resistance. Our results are consistent with reports from other groups. There is evidence that polystyrene (PS) microplastics increase the risk of insulin resistance in mice by affecting the gut-liver axis (Shi et al., 2022). Another study in mice showed insulin resistance after PS exposure in association with increased plasma proinflammatory cytokines such as tumor necrosis factor-α and interleukin-1β (Huang et al., 2022). Polystyrene microplastics also elevated ROS in the liver, which disrupted the PI3K/Akt signaling pathway and led to insulin resistance (Fan et al., 2023, 2022). It is worth noting that our previous study demonstrated the effects of PET microplastics on the expression of porcine serum-derived extracellular vesicles miRNA. These differentially regulated miRNAs can be involved in the development of metabolic syndrome, insulin resistance and type 2 diabetes (Mierzejewski et al., 2023). There is evidence that insulin resistance is associated with a high rate of cholesterol synthesis and low cholesterol absorption (Pihlajamäki et al., 2004). Interestingly, our study showed a stimulatory effect of a low dose of PET microplastics on serum cholesterol levels, supporting the assumption that PET microplastics may significantly disrupt insulin homeostasis.

An important element of our research was the finding of a stimulatory effect of a low dose of PET microplastics on γ-aminobutyric acid (GABA) levels in the pancreas. It is known that pancreatic β-cells synthesize GABA from glutamic acid by GAD and store it in the synaptic macrovesicles with insulin granules until the time of secretion (Al-Kuraishy et al., 2021). It has been reported that GABA plays a fundamental role in the regulation of pancreatic β-cells by suppressing and stimulating glucagon and insulin secretion, respectively. In addition, GABA has the specific ability to restore β-cells after their destruction (Wang et al., 2019). This phenomenon could be a compensatory mechanism of the pancreas in response to the increased glucose levels in pancreatic cells under the influence of microplastics. However, the significance of this mechanism requires clarification in subsequent studies.

An important group of metabolites affected by PET microplastics in the pancreas are lysophospholipids such as lysophosphatidylcholine (lysoPC, LPC) and lysophosphatidylethanolamine (lysoPE, LPE). They are bioactive lipids with diverse physiological and pathological functions in the pancreas. LPC is known as an endogenous mediator that triggers insulin secretion in pancreatic β-cells. LPC has been reported to increase the transcriptional activity of NF-kB and AP-1 and induce apoptosis by activating extracellular signal-regulated kinase (ERK) 1/2, c-Jun NH2-terminal kinase/stress-activated protein kinase (JNK/SAPK) and p38 MAP kinases in rat pancreatic AR42J cells (Masamune et al., 2001). The prospective study using prediagnostic blood samples showed that higher LPC and LPE levels were associated with the risk of pancreatic ductal carcinoma (Naudin et al., 2023). The present study showed that a low dose of PET microplastics upregulated the levels of LPC (18:2), LPC (20:3), LPC (20:2) and LPC (22:5) in the pancreas. In turn, a high dose of PET microplastics increased LPE (20:1) in the pancreas. The results suggest that exposure to PET microplastics, even at low doses, can significantly alter lipid metabolism in the pancreas, potentially increasing the risk of pancreatic diseases, including pancreatitis, or the risk of pancreatic cancer. Another metabolite that may serve as a marker for chronic pancreatitis is glycerophosphocholine (GPC), which is catalyzed from LPC (Lu, 2012). Numerous studies have indicated that choline metabolism is altered in a variety of cancers. Activated choline metabolism is a hallmark of carcinogenesis and leads to elevated phosphocholine and glycerophosphocholine levels in multiple cancers (Sonkar et al., 2019). Our studies have shown an increase in GPC levels in the pancreas after exposure to low doses of PET microplastics.

We found that PET microplastics increased the level of pancreatic lipase in serum fourfold compared to the control. The measurement of pancreatic lipase levels is routinely used in clinical practice for the diagnosis of pancreatic diseases. Their elevated levels correlate strongly with various pathological conditions, especially acute and chronic pancreatitis (Smith et al., 2005). Moreover, there is evidence that persistently elevated or increasing combined levels of amylase and lipase in serum are reliable indicators of pancreatic injury (Mahajan et al., 2014). Although our studies showed no significant change in serum amylase levels due to PET exposure, given the significantly elevated lipase levels and the literature data indicating that microplastics cause pancreatic injury in mice (Zheng et al., 2023), we suggest that future research should consider assessing the extent of tissue damage over time and the levels of pancreatic enzymes in serum. Furthermore, our study revealed that PET microplastics in high doses increase serum calcium levels. It is well documented that abnormal regulation of Ca^2+^ signaling is one of the central triggers for the pathogenesis of acute pancreatitis (Mounzer et al., 2012). Hypocalcemia also was significantly more frequent in patients with a severe form of acute pancreatitis than in patients with a mild form (Mounzer et al., 2012).

In summary, the etiology and pathogenesis of various pancreatic diseases such as pancreatitis, pancreatic cancer or insulin resistance remains unclear. In many diseases, metabolic abnormalities occur before changes in tissue structure and functional changes. Therefore, a global metabolomic approach is extremely important to identify early pathological changes that may lead to disease. This study is the first to provide a comprehensive overview of the effects of PET microplastics on pancreatic function. Our research has shown that microplastics, both at low and high doses, affect the physiological processes of the pancreas in young organisms. We postulate that PET microplastics increase the risk of insulin resistance, pancreatitis and pancreatic cancer.

## Author contributions

Conceptualization: K.M., I.B.; Data curation: K.M.; Formal analysis: K.M.; Funding acquisition: K.M.; Investigation: K.M., A.K., M.G. I.G., J.C.; Methodology: K.M., I.G. J.C.; Project administration: K.M.; Resources: K.M., A.K., M.G.; Software: K.M.; Supervision: I.B.; Visualization: K.M. I.G.; Writing: original draft: K.M.; Writing: review and editing: K.M., I.B., All authors have read and agreed to the published version of the manuscript.

## Acknowledgement

We would like to thank to MSc Karolina Łowicka for technical assistance during performing the experiments.

## Data availability statement

The raw data generated for this study can be found in the EMBL-EBI MetaboLights database with the identifier MTBLS10732.

## Supplementary materials

**Table S1** (a) List of peak intensities identified in the study after normalization against QC samples in negative ionization mode (ESI-) after treatment of the piglets with a low dose of PET microplastics. (b) List of statistically significantly different metabolites between control samples and those treated with a low dose of PET microplastics, identified in the study in negative ionization mode (ESI-). (c) KEGG pathways enrichment analysis of metabolites identified in the study in negative ionization mode (ESI-).

**Table S2** (a) List of peak intensities identified in the study after normalization against QC samples in positive ionization mode (ESI+) after treatment of the piglets with a low dose of PET microplastics. (b) List of statistically significantly different metabolites between control samples and those treated with a low dose of PET microplastics, identified in the study in positive ionization mode (ESI+). (c) KEGG pathways enrichment analysis of metabolites identified in the study in positive ionization mode (ESI+).

**Table S3** (a) List of peak intensities identified in the study after normalization against QC samples in negative ionization mode (ESI-) after treatment of the piglets with a high dose of PET microplastics. (b) List of statistically significantly different metabolites between control samples and those treated with a high dose of PET microplastics, identified in the study in negative ionization mode (ESI-). (c) KEGG pathways enrichment analysis of metabolites identified in the study in negative ionization mode (ESI-).

**Table S4** (a) List of peak intensities identified in the study after normalization against QC samples in positive ionization mode (ESI+) after treatment of the piglets with a high dose of PET microplastics. (b) List of statistically significantly different metabolites between control samples and those treated with a high dose of PET microplastics, identified in the study in positive ionization mode (ESI+). (c) KEGG pathways enrichment analysis of metabolites identified in the study in positive ionization mode (ESI+).

